# Coordination of robust single cell rhythms in the *Arabidopsis* circadian clock via spatial waves of gene expression

**DOI:** 10.1101/208900

**Authors:** Peter D. Gould, Mirela Domijan, Mark Greenwood, Isao T. Tokuda, Hannah Rees, Laszlo Kozma-Bognar, Anthony J.W. Hall, James C.W. Locke

**Affiliations:** Institute of Integrative Biology, University of Liverpool, Crown Street, Liverpool, L69 7ZB, UK; Sainsbury Laboratory, University of Cambridge, Bateman Street, Cambridge, CB2 1LR, UK; Department of Mathematical Sciences, University of Liverpool, Peach Street, Liverpool, L69 7ZL; Department of Biochemistry, University of Cambridge, 80 Tennis Court Road, Cambridge, CB2 1QW, UK; Department of Mechanical Engineering, Ritsumeikan University, Noji-higashi, Kusatsu, 5258577, Japan; Biological Research Centre, Hungarian Academy of Sciences, 6726 Szeged, Hungary; Earlham Institute, Norwich Research Park, Norwich, NR4 7UH, UK; Microsoft Research, 21 Station Road, Cambridge, CB1 2FB, UK

## Abstract

The *Arabidopsis* circadian clock orchestrates gene regulation across the day/night cycle. Although a multiple feedback loop circuit has been shown to generate the 24h rhythm, it remains unclear how robust the clock is in individual cells, or how clock timing is coordinated across the plant. Here we examine clock activity at the single cell level across *Arabidopsis* seedlings over several days. Our data reveal robust single cell oscillations, albeit desynchronised. In particular, we observe two waves of clock activity; one going down, and one up the root. We also find evidence of cell-to-cell coupling of the clock, especially in the root tip. A simple model shows that cell-to-cell coupling and our measured period differences between cells can generate the observed waves. Our results reveal the spatial structure of the plant circadian clock and suggest that unlike the centralised mammalian clock, the clock has multiple points of coordination in *Arabidopsis*.

## Introduction

The circadian clock controls gene expression throughout the day and night in most organisms, from single cell photosynthetic bacteria to mammals (Bell-Pedersen et al., 2005). In many cases a core circuit that generates this rhythm has been elucidated and been shown to oscillate in single cells. In multi-cellular organisms these single cell rhythms must be integrated to allow a coordinated response to the environment. Mammals achieve this by driving oscillations in peripheral tissues from a central pacemaker in the brain, the suprachiasmatic nucleus (SCN) (Pando, Morse, Cermakian, & Sassone-Corsi, 2002; Reppert & Weaver, 2002).

The *Arabidopsis* circadian clock generates a 24h rhythm in multiple key processes, including stomata opening, photosynthesis, and hypocotyl elongation (Hsu & Harmer, 2014). A hierarchical structure for the plant clock has recently been proposed, similar to that for the mammalian clock, where the shoot clock drives the rhythms in the leaves and roots (Takahashi, Hirata, Aihara, & Mas, 2015). However, there are further tissue dependent differences that must be explained. For example, experiments using a luciferase reporter for clock activity have shown waves of clock gene expression in leaves (Fukuda, Nakamichi, Hisatsune, Murase, & Mizuno, 2007; Wenden, Toner, Hodge, Grima, & Millar, 2012), as well as striped expression patterns in roots (Fukuda, Ukai, & Oyama, 2012).

Beyond the coordination of plant rhythms, how robust the circadian clock is in individual cells across the plant is also unclear. Through integration of data from whole plant studies, a genetic circuit consisting of multiple coupled feedback loops has been proposed to generate the 24h rhythm (Fogelmark & Troein, 2014; Pokhilko et al., 2012). Simulations of this network display stable oscillations (Figure 1a), although experimental measurements of clock rhythms under constant conditions often display damped rhythms (Figure 1b) (Gould et al., 2013; Locke et al., 2005, 2006; Salomé & McClung, 2005). This damping could be due to the clock circuit in individual cells losing rhythmicity (top, Figure 1c), or to cells desynchronising due to different intrinsic periods or phases (Guerriero et al., 2012; Komin, Murza, Hernández-García, & Toral, 2010) (bottom, Figure 1c), or cells desynchronising due to stochasticity in clock activity (Guerriero et al., 2012). Previous studies have attempted to measure the clock in plants at single-cell resolution; however, these have been confounded by poor temporal/spatial resolution and short time series (Takahashi et al., 2015; Yakir et al., 2011).

**Figure 1.**
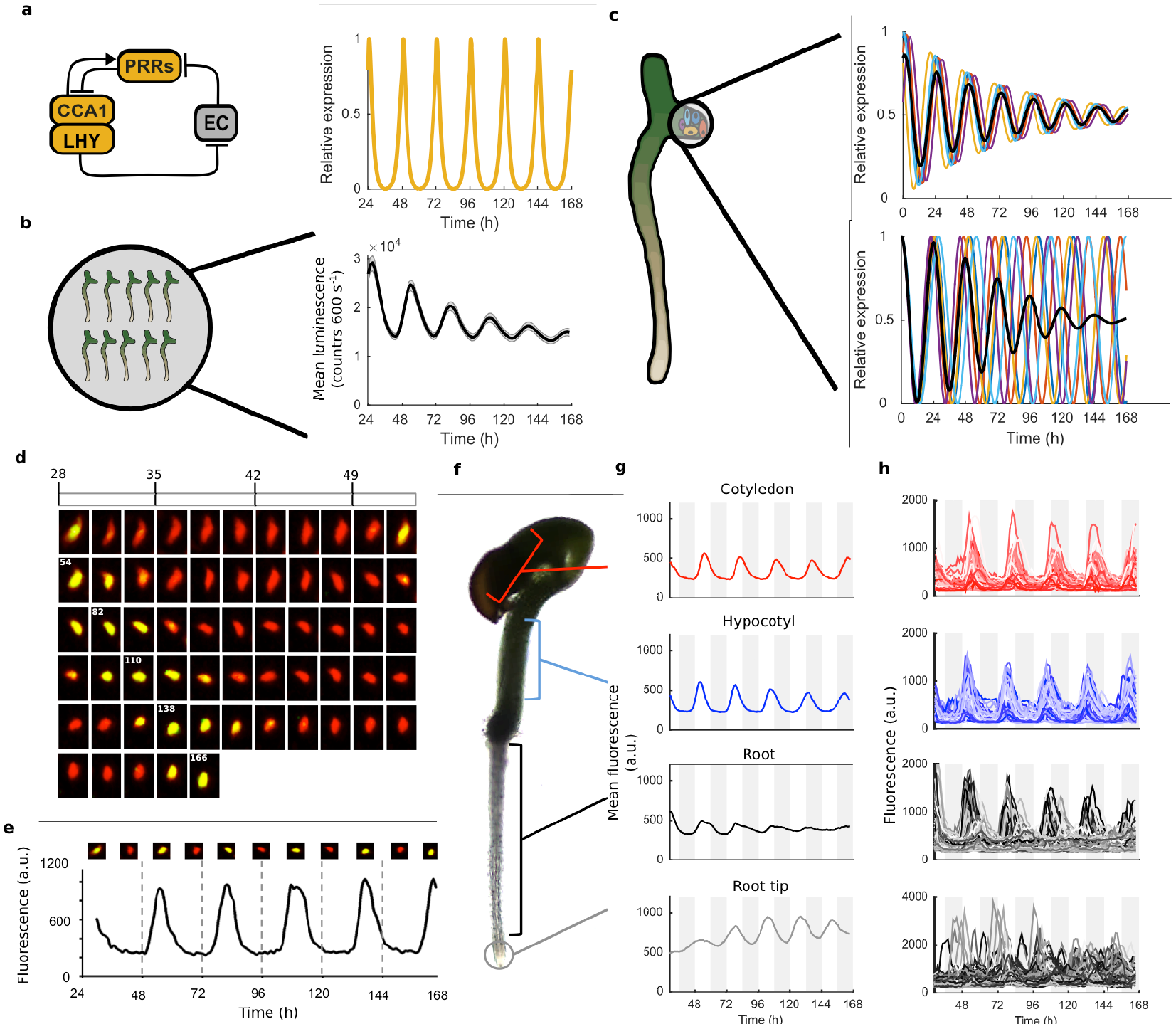
Quantitative time-lapse microscopy reveals single cell clock dynamics across the plant. **a**, Current models of the clock (Pokhilko et al., 2012) predict undamped oscillations, **b**, CCA1:LUC bulk averaged over multiple seedlings shows damping clock oscillations (mean ± s.e.m.; *n* = 32 seedlings), **c**, The reason for damping could be due to damping rhythms in individual cells (top) or due to desynchronisation between cells (bottom), **d**, Images show a representative nuclei from the cotyledon of a *cca1-11* CCA1::CCA1-YFP; 35S::H2B-RFP seedling that had been grown in LD cycles before being released into continuous light free running conditions for several days. The red channel represents when H2B fluoresce and the yellow CCA1. Times of peak expression are indicated on images. **e**, Expression levels of CCA1-YFP from the representative nuclei shown in (**d**). Images of the nuclei are also shown for the peaks and troughs in the CCA1-YFP oscillation. **f**, Representative seedling identifying the different sections imaged. **g**, Mean traces of single cell CCA1-YFP for over 5 days of constant light in different regions of the plant showing damping rhythms in the root, but not the root tip. **h**, CCA1-YFP traces from individual cell in each section for the same 5 days.

Here, we examine the dynamics of the *Arabidopsis* clock across the whole plant at the single cell level over several days. Our results reveal that damping of rhythms is mainly due to desynchronisation of oscillating single cells with different periods, and not due to noise in gene expression or lack of robustness. We observe two waves of clock gene expression, one up and one down the root, which cause the most desynchronisation. From our single cell data, we are able to estimate the coupling strength between cells, and find evidence of coupling, especially strong in the root tip. A simple model suggests that our observed period differences, plus cell-to-cell coupling, can generate the observed waves in clock gene expression. Thus, our data has revealed both the structure and robustness of the plant circadian clock system.

## Results

To analyse the dynamics of the plant clock at the single cell level, we constructed reporter lines that allowed us to quantitatively measure the nuclear level of the core clock protein CIRCADIAN CLOCK ASSOCIATED 1 (CCA1) (Wang & Tobin, 1998). These reporter lines contained a CCA1-YFP protein fusion construct driven by the *CCA1* promoter in a *cca1-11* mutant background. They also contained a 35S::H2B-RFP nuclear marker to enable automatic detection of individual nuclei (Federici, Dupuy, Laplaze, Heisler, & Haseloff, 2012). By screening the clock phenotypes of multiple reporter lines we ensured that our reporter construct was functional and rescued the period phenotype of the *cca1-11* mutant (Figure 1-figure supplement 1). We took forward both a rescued wild-type period (WT) and a long period (CCA1-long) reporter line for further analysis.

We carried out time-lapse movies of *Arabidopsis* seedlings using a custom developed time-lapse confocal microscope setup (Figure 1-figure supplement 2). In order to examine the intrinsic behavior of the clock we first entrained the seedlings to 12:12h light/dark cycles before examining the clock under constant conditions (constant blue light (30 μmol m^−2^ s^−1^) and temperature (22 °C)), as standard in circadian research. Our method allowed us to track and extract fluorescence values from the same individual nuclei over several days (Figure 1d, e). We first examined the average CCA1-YFP nuclear fluorescence signal from regions of the hypocotyl, cotyledon and roots (Figure 1f, g). We observed a robust oscillation in the cotyledon (red line, Figure 1g) and hypocotyl (blue line, Figure 1g), although with slight damping to the amplitude. In the top part of the root we observed strong damping of the circadian rhythm (black line, Figure 1g), although surprisingly the oscillations recovered somewhat in the root tip (grey line, Figure 1g). Three repeat plants showed similar behavior (Figure 1-figure supplement 3). We also observed similar behavior in our CCA1-long reporter line (Figure 1-figure supplement 4), showing that our results remain true across a range of clock activity.

To determine what the underlying cause of the damping in different tissues was we examined the clock rhythm in thousands of individual cells (Figure 1h, Figure 1-figure supplement 3b, 4b, 5). Oscillations could be observed in all tissues, with most cells displaying circadian oscillations (Figure 1, Figure 1-source data 1, 2). It is clear from the traces that the strong damping in the mean levels in the root is not caused by individual oscillators losing rhythmicity (Figure 1h). We then examined the synchronicity and robustness of these rhythms in more detail. The hypocotyl and cotyledon were the most synchronised, with the amplitude of the mean trace nearly equaling the median amplitude of the individual cell lineages (Figure 2a, Figure 2-figure supplement 1, 2). The hypocotyl and cotyledon rhythms also exhibited low period variability both within and between cell lineages, indicating a high level of robustness (Figure 2b, Figure 2-figure supplement 1b, 2b). However, the root displayed significant desynchronisation, with the amplitude of the mean trace lower than the median amplitude of the individual cell lineages (Figure 2a, Figure 2-figure supplement 1a, 2a), with higher variability in period within and between single cell lineages (Figure 2b and Figure 2-figure supplement 1b, 2b).

**Figure 2.**
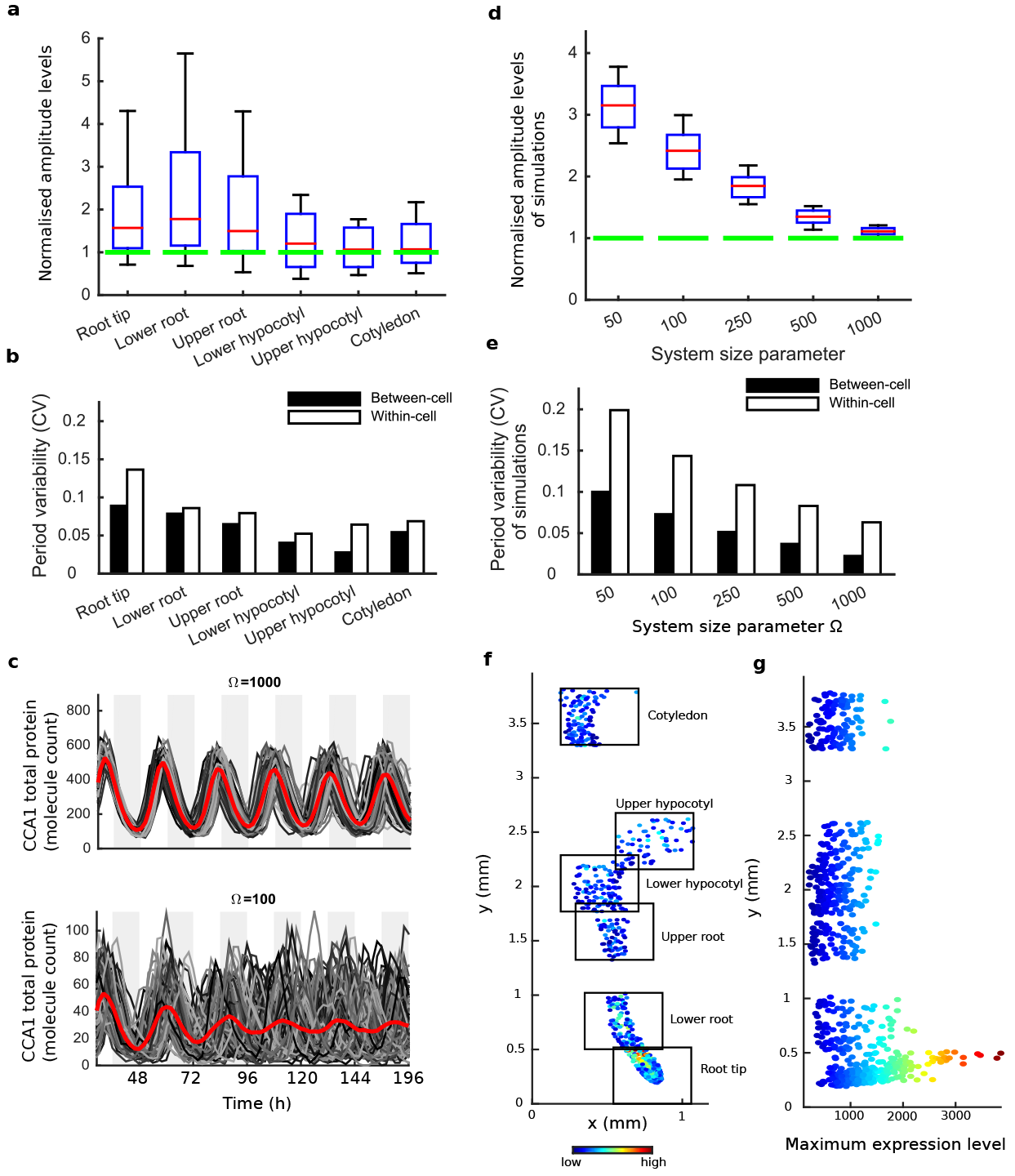
Single cell analysis reveals tissue level differences in robustness of the clock. **a**, Rhythmic cell amplitudes in the imaged sections normalised to the amplitude of the mean trace (green line). For root tip, *n* = 242; lower root, *n* = 83; upper root, *n* = 46; lower hypocotyl, *n* = 114; upper hypocotyl, *n* = 53; cotyledon, *n* = 103. Whiskers represent 9^th^ and 91^st^ percentile, *n* the number of cells. **b**, Between-cell and within-cell period variability in each imaged section. **c**, Stochastic model CCA1 total molecule count for Ω = 1000 (top) and Ω = 100 for 100 (bottom) simulated runs (grey) plotted from 29h to 168h in constant light (comparable to the data in Figure 1). Means of all simulated runs are shown in red. Ω represents the system size. **d**, Rhythmic simulated run amplitudes for different system sizes (Ω) normalised to the mean simulation (green line). **e**, Between and within cell variability of each simulation with different scaling factor. **f**, Scatterplot of the rhythmic cells in all imaged plant sections stitched together. Colour indicates the oscillation amplitude. **g**, Scatterplot of the amplitude values vs. longitudinal position on the plant measured from the root tip. Colour legend is the same as (**f**).

We next asked whether the damping of amplitude in the mean rhythm is caused by noise in gene expression, as proposed by previous stochastic modelling (Guerriero et al., 2012). Simulations of the stochastic model of the *Arabidopsis* gene network (Guerriero et al., 2012) display greater desynchronisation as the system size, effectively the simulated number of molecules in the cell, is reduced (Figure 2c-e). The lower the molecule number, the more desynchronisation at the single cell level, and the more damping of the mean expression (red line, Figure 2c). To explore this further, we simulated the model for a range of system sizes (Figure 2c) and examined the synchronicity and robustness of the simulations (Figure 2d, e). Previously the system size was estimated to be of order 100 molecules per cell (bottom panel, Figure 2c) in order for the desynchronisation observed in whole plant measurements to be explained solely by noise (Guerriero et al., 2012). We compared the level of desynchronisation and noise from simulations of this model to our experimentally measured single cell rhythms. The single cell rhythms that we detect in the hypocotyl and cotyledon are more robust than a system size of 100, and in fact behave closer to a system size of 1000 (top panel, Figure 2c). The level of desynchronisation observed in the root and root tip, however, behave more like a simulated system size of 100 (Figure 2b, c). Similar CCA1-YFP expression levels are observed across the plant (Figure 2f, g), so it is unlikely that this lack of robustness is due solely to a change of total molecule number for the clock system in different parts of the plant. In fact the amplitude and expression levels of the single cell oscillations in the root tip are high compared to other sections across the plant (Figure 2f, g and Figure 2-figure supplement 1a, c-d, 2a, c-d).

If the damping in the root is not due to noise in gene expression, or individual cells losing rhythmicity, what does cause it? The measured variable and desynchronous rhythms in the root suggest that the clock could be behaving differently in different parts of the root. To test this possibility we plotted the period of the individual cell oscillations across the plant (Figure 3a, b and Figure 2-figure supplement 1e, f and 2e, f). We observed surprising spatial structure to the clock in the root. The upper sections of the root displayed longer periods than the rest of the plant, as reported previously for the whole root (James et al., 2008). However, we observed very fast rhythms in the root tip (Figure 3a, b), which is also the section with very high expression rhythms (Figure 2f, g). Although we do not observe evidence of phase resetting in the root tip, as proposed in an earlier luciferase study (Fukuda et al., 2012), this could be due to our different growth conditions and stage of plant development. A repeat plant showed similar results (Figure 2-figure supplement 1c-f), as did the CCA1-long reporter line (Figure 2-figure supplement 2c-f). Each section of the plant can be made up of multiple cell types. So, we next tested whether the rhythms have any spatial structure in the *z* direction, which would suggest that different cell types have different period rhythms. Plots of period in the *z* direction in each section do not reveal any discernable pattern, including in the root, where cells are organised radially (Figure 3c, d). This suggests that the differences in rhythms we observe are not restricted to a specific cell type.

**Figure 3.**
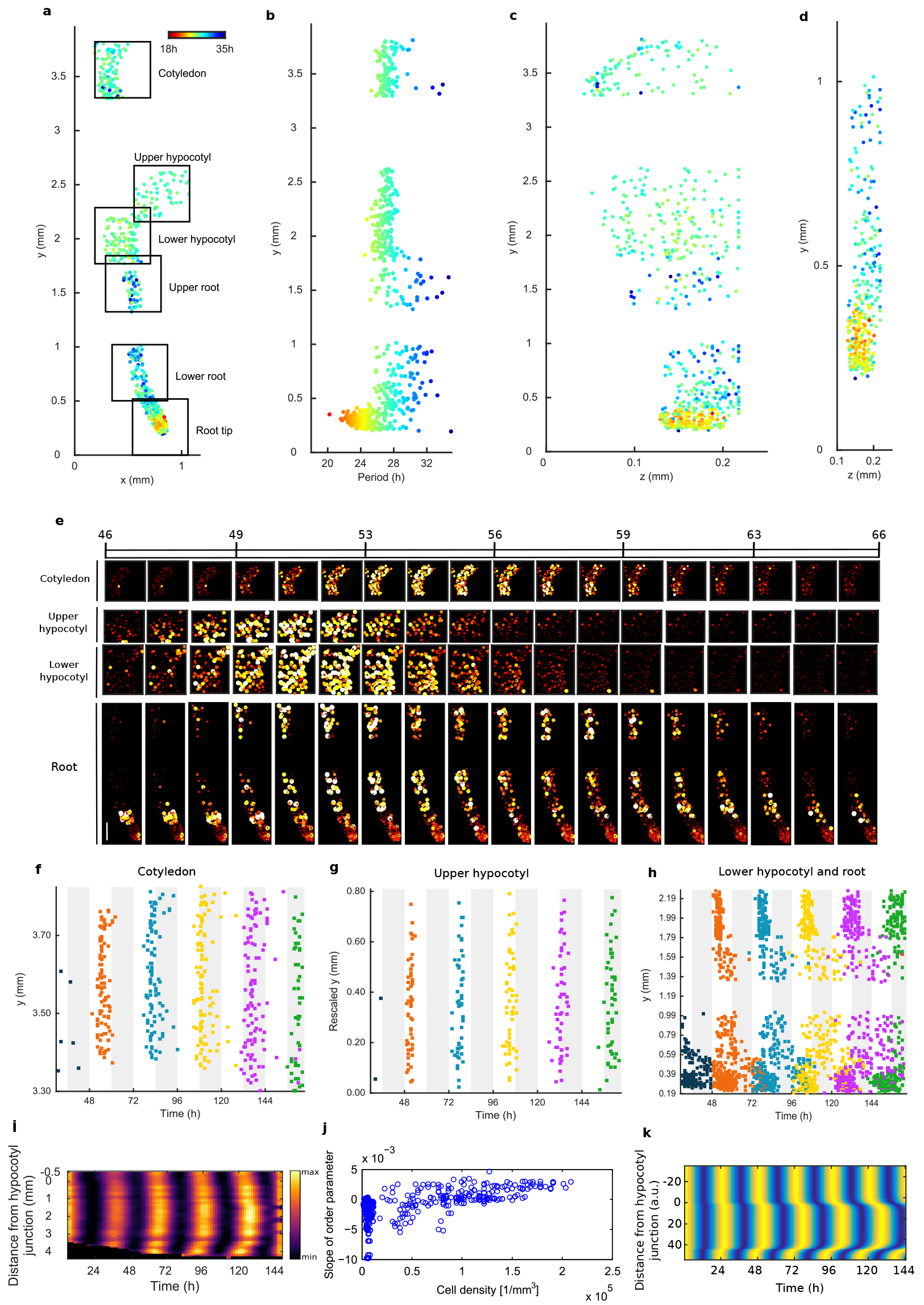
Single cell period differences and cell-to-cell coupling generate spatial waves of clock gene expression. **a**, Scatterplot of the rhythmic cells in all imaged plant sections stitched together in *x*-*y* direction. Colour indicates the oscillation period. **b**, Period values vs. longitudinal position on the plant measured from the root tip. Colour legend is the same as (**a**). **c, d**, Scatterplot in the *y*-*z* direction of rhythmic cells in all imaged plant sections (**c**) or in the root and root tip sections only (**d**). Colour legend is the same as (**a**). **e**, Montage of the normalised expression of rhythmic cells from the root (bottom panel, first image taken after 46.1h in LL), lower hypocotyl (taken after 46.6h in LL), upper hypocotyl (taken after 46.7h) and cotyledon (top panel, taken after 46.9h in LL). Each frame is approximately 1.1h apart. Scale bar represents 0.25 mm. **f-h**, Space-time plots of peak times of rhythmic cells across sections: cotyledon (**f**) upper hypocotyl (**g**), lower hypocotyl and root (all sections) (**h**). **i**, Representative space-time plot of normalised PRR9:LUC expression across longitudinal sections of a seedling (N = 2, *n* = 7). *N* represents the number of independent experiments, *n* the total number of individual seedlings. **j**, Slopes of the order parameter are plotted against cell densities. A positive slope indicates that the level of synchrony increases in time due to cell-cell interactions. **k**, Space-time plot of simulated total normalized *CCA1* expression across longitudinal sections of the seeding.

To further understand the spatial structure of the clock, we examined montages of clock gene expression (Figure 3e; Figure 2-figure supplement 1g; Videos 1-3) and timing of peaks of expression (Figure 3f-h) across the plant. The clock peaks earlier in the hypocotyl than in the cotyledon (Figure 3f-h), and the phase of the rhythm is also tightest in the hypocotyl, where the phase varies by a standard deviation of 1.47h in the first peak and 3.09h in last peak (Figure 3g), with more desynchronisation in the cotyledon (Figure 3f). From the top of the root the phase of the clock is shifted to later in the day as you go down the root (Figure 3h). However, from the root tip the phase of the clock is shifted to later in the day as you go up. This generates two waves in the montages of clock gene expression, one going up and one going down the root (Figure 3e; Video 1; Figure 2-figure supplement 1g, 2g). Thus, the damped mean rhythm of clock activity we observed in the root (Figure 1g) is caused by the averaging of these two waves of gene expression. We observed qualitatively similar waves in two clock luciferase reporter lines, PSEUDO-RESPONSE REGULATOR 9:LUC (Salomé & McClung, 2005) (Figure 3i; Video 4) and CCA1:LUC (Figure 3-figure supplement 1), showing that our results are a general property of the clock.

The coherent waves of gene expression suggested that the plant clock signal could be coupled. To estimate this coupling, we calculated the order parameter (Kuramoto, 1984) from our single cell data and estimated the coupling strength based on a technique developed for mammalian circadian cells (Rougemont & Naef, 2007). We observed signs of coupling across the plant, with the strongest evidence for coupling in the root tip, where the order parameter actually increased with time (Figure 3j and Figure 2-figure supplement 2-4). Interestingly this occurred where cell density was highest, as observed in cultured SCN cells (Aton, Colwell, Harmar, Waschek, & Herzog, 2005). To investigate the mechanism for the waves of clock gene expression in the root we developed a simple mathematical model where the cells are described by coupled phase oscillators with different periods, as informed by the data (Figure 3k). The periods of the clock in the model were faster in the shoot and root tip than the rest of the root, as measured experimentally. This simple model can generate waves of gene expression up and down the root that produce a bow wave in the space-time plot (Figure 3k), similar to that observed experimentally (Figure 3h, i).

## Discussion

Our single cell measurements have revealed tissue specific differences in the phases and robustness of the clock in *Arabidopsis*. These differences are not restricted to one cell type, as similar periods are observed in the *z* dimension through the plant (Figure 3c, d), suggesting that cells are instead responding to information based on their longitudinal position. The observed robust rhythms in the hypocotyl that peak before the cotyledon and roots are in line with a proposed hierarchical structure for the plant clock, where the shoot clock drives the rhythms in the leaves and roots (Takahashi et al., 2015). However, our results suggest that the structure of the plant clock is more complicated, as this hierarchical model does not explain the observed short period oscillations in the root tip. Our results support a more decentralised model of clock coordination in plants (Endo, 2016; Endo, Shimizu, Nohales, Araki, & Kay, 2014).

Earlier studies of the clock argue either that the clock is uncoupled (Thain, Hall, & Millar, 2000; Yakir et al., 2011), or weakly, but detectably, coupled (Fukuda et al., 2007, 2012; James et al., 2008; Takahashi et al., 2015; Wenden et al., 2012). Our single cell approach is consistent with weak coupling across the whole plant but reveals regions with strong local coupling between cells, especially in the root tip, which is sufficient to drive an increase in synchrony with time. Our modelling shows that this coupling together with the observed period differences is sufficient to replicate the decentralised spatial structure of the clock that we observe experimentally.

Decentralised coordination could create flexibility and allow parts of the plant to respond differentially to environmental perturbations. There is already evidence that the root clock may respond differently to light (Bordage, Sullivan, Laird, Millar, & Nimmo, 2016), and that the vasculature and epidermal clock regulate distinct physiological processes (Shimizu et al., 2015). It has been recently shown that initiation of lateral roots triggers the resetting of the clock in the emerging lateral root (Voß et al., 2015). In the case of lateral roots auxin is proposed to be involved in resetting the clock (Voß et al., 2015). An important next step will be to investigate what the coupling signal is for the plant circadian clock (Covington & Harmer, 2007; Dalchau et al., 2011; Haydon, Mielczarek, Robertson, Hubbard, & Webb, 2013).

## Acknowledgements

We would like to acknowledge the Liverpool Centre for Cell Imaging for assistance and maintenance of the confocal microscope, specifically Marco Marcello facility manager and David Mason image analysis support. A.H. and P.D.G. were funded by BBSRC grant BB/K018078/1, J.C.W.L. and M.D. by BBSRC grant BB/K017152/1 and M.G. by the Liverpool/Durham/Newcastle BBSRC DTP. Correspondence and requests for materials should be addressed to J.L. or A.H.

## Contributions

A.J.W.H. and J.C.W.L. designed the research. P.D.G. developed the time-lapse microscopy platform and carried out the microscopy and delayed fluorescence imaging. M.D. carried out the mathematical modelling and statistics. M.G. carried out the luciferase imaging and analysis. L.K. completed the cloning. P.D.G., H.R., M.D., M.G. and I.T. carried out analysis of single cell data. All authors contributed to the writing of the manuscript.

## Competing interests

The authors declare no competing financial interests.

## Methods

### Constructs

The CCA1∷CCA1-YFP was constructed as follows: The coding region of *CCA1* was amplified from *Arabidopsis* (Ws) genomic DNA using the primer pair CCA1_CDS_Fwd (5’-AAAGGATCCATGGAGACAAATTCGTCTGGA-3’) and CCA1_CDS_Rev (5’-ATACCCGGGTGTGGAAGCTTGAGTTTCCAA-3’). The *CCA1* promoter region was amplified from *Arabidopsis* (Ws) genomic DNA using the primer pair CCA1_prom_Fwd (5’-AAAGAATTCATTTAGTCTTCTACCCTTCATGC-3’) and the CCA1_prom_Rev (5’-ATAGGATCCCACTAAGCTCCTCTACACAACTTC-3’). Unique restriction sites were designed at the ends of the amplicons to facilitate cloning. The fragment of *CCA1* coding region was cloned in the modified pPCV812 binary plasmid(Pfeiffer et al., 2009) between the *35S* promoter and the *YFP* gene via *Bam*HI (5’) and *Sma*I (3’) sites, resulting in 35S∷CCA1-YFP. Next, the *35S* promoter was replaced by the *CCA1* promoter fragment via *Eco*RI (5’) and *Bam*HI (3’) sites, resulting in CCA1∷CCA1-YFP. The cloned *CCA1* promoter fragment was 862 bp in length and contained the full 5’ -untranslated region, but not the ATG.

### Plant growth material

The CCA1∷CCA1-YFP construct was transformed in *cca1-11* (Ws) mutant background (Hall et al., 2003). Homozygous T3 generations of several independent transgenic lines were checked for complementation via delayed fluorescence (Gould et al., 2009) (Figure 1-figure supplement 1b, c). The CCA1∷CCA1-YFP expressing line showing full complementation was then re-transformed with *35S*∷H2B-RFP which was used for tracking purposes during analysis (Federici et al., 2012). The CCA1:LUC and PRR9:LUC lines used are in the Col-0 background and were built as part of the ROBuST project.

*Arabidopsis* seed were surface sterilised and suspended in 0.1% top agar and placed in 4 °C for 2 days. After sowing, seeds were grown inside of growth incubators (Sanyo MLR-352) under 12:12 LD cycles in 80 umol m^2^ s^−1^ cool white light at 22 °C for entrainment. Henceforth, these conditions are referred to as entrainment conditions.

### Luciferase and delayed fluorescence bulk imaging

10-20 seed were sown onto Murashige and Skoog (MS) 2% agar in clear 96 well microtitre plates with at least 8 wells per line. A second clear microtitre plate was placed on top of the plate containing the seed to increase well height. These were then sealed using porous tape (Micropore). Seedlings were grown for 9 days under entrainment conditions and transferred to experimental conditions on dawn of the 10^th^ day. For luciferase experiments, on the 9^th^ day seedlings were sprayed with a 5 mM luciferin solution in 0.001% Triton x-100 before transfer to experimental conditions on dawn of the 10^th^ day.

Imaging was carried out in Sanyo temperature controlled cabinets (MIR-553 or MIR-154) at 22 °C and under an equal mix of red and blue LEDs (40 μmol m^−2^ sec^−1^ total). Seedlings were imaged using an ORCA-II-BT (Hamamatsu Photonics, Japan) or LUMO CCD camera (QImaging, Canada). Experiments were run over several days with images being taken every hour as described previously (Gould et al., 2013; Litthauer, Battle, Lawson, & Jones, 2015). Image analysis was carried out using Imaris (Bitplane, Switzerland) or ImageJ (NIH, USA).

### Luciferase macro imaging

Seedlings were sown in a row of eight seedlings on 2% agar media supplemented with MS media. Seedlings were grown upright for four days under entrainment conditions. Imaging commenced on dawn of the fifth day. Imaging was performed inside of Sanyo plant growth incubators at 22 °C under an equal mix of red and blue light emitting diodes (40 μmol m^−2^ sec^−1^ total). Seedlings were imaged upright using a LUMO CCD camera (QImaging, Canada). Experiments were run over several days with images being taken every 90 minutes.

### Confocal microscopy

For confocal experiments seed were sown directly onto glass bottom dishes (Greiner, Austria) in an array format. Once dry, the seed were covered with 5 ml MS 2% agar media in absence of sucrose. Once set, dishes were sealed with porous tape (Micropore) and grown upright under entrainment conditions for 4 days. After 4 days plates were ready for imaging.

The microscopy pipeline is outlined in Figure 1-figure supplement 2. Up to 18 seedlings were grown in an array format on glass bottom dishes. At dawn on the 4^th^ day of growth the dishes were fixed into the confocal temperature controlled stage (22 °C) using the dish manifold. To maintain correct light conditions (30 μmol m^−2^ s^−2^ constant blue light) a custom-made light emitting diode (LED) rig was used. Growth conditions allowed slow growth during the movie, which enabled easier tracking of single cells. A Zeiss 710 (Zeiss, Germany) inverted confocal microscope with a 40x/1.2 water corrected oil objective was used for all imaging. YFP and RFP excitation was produced using a 514 nm laser and a main beamsplitter (MBS) 458/514. To reduce problems with auto-fluorescence and improve signal to noise ratio a lambda scan was carried using a ChS PMT and filters 492-658. Brightfield (BF) used a ChD PMT. Imaging was carried out using a 0.6 zoom to increase field of view. A motorised stage was used to allow multiple positions to be imaged across the plant per experimental run. The diameter of nuclei in our seedlings ranges in size from 6 μm (root tip) to 15 μm (hypocotyl). A resolution of 2 μm in the *z* dimension was chosen to allow the capture of several slices through each nucleus. Data was auto saved during imaging with data split into files by position imaged.

### Processing confocal images

Firstly blank images created by time-lapse being terminated early were removed in ImageJ. Also with ImageJ, lambda scans produced during confocal imaging were split into YFP (511 to 547 nm), RFP (586 to 625 nm) and brightfield (BF) spectrums and then reduced in dimensionality to give one channel for each wavelength. Data was then saved as OME TIFF, writing each time point as a separate file. Once processed all the data was loaded into Imaris (Bitplane, Switzerland) and merged to produce one file containing YFP, RFP and BF. A median filter size 3x3x1 was then applied across all of the data. Detection of YFP/RFP expressing cells was carried out using the spot detection feature and tracking of spots over time was carried out using an autoregressive motion model using an estimated cell *x,y* diameter of 6-10 μm. Data was then exported in excel format for further analysis. Details are provided in the subsequent section. Quality control checks were carried out at multiple points (see Figure 1-figure supplement 2). The first quality check was made to ensure that the seedling remained in the focal plane during the course of the experiment. If not, the dataset was not carried forward for further analysis. The second check was to make sure that all processing has occurred correctly. The third check was carried out to correct any errors in tracking cells across the time-lapse data. The fourth check used the videos to more closely monitor the data for anything that looked problematic. The final check used the graphs to identify any problems that may have occurred during the whole single cell pipeline. If the laser power was not found to be stable during the course of the imaging the dataset was not carried forward for further analysis.

### Single cell data processing

Period analysis was carried out in BioDare, an online system for data sharing and analysis (Costa et al., 2013; Moore, Zielinski, & Millar, 2014). Since most of the period analysis methods in BioDare require evenly spaced time series, the data was first interpolated (using MATLAB’s (MathWorks, U.K.) interp1 function and spacing of 1h). Period estimates were obtained by three different methods: Spectrum Resampling (Costa et al., 2013), FFT-NLLS (Johnson & Frasier, 1985; Straume, Frasier-Cadoret, & Johnson, 2002) and mFourFit (Edwards et al., 2010). Cells were classed as rhythmic only if each method identified them as rhythmic (i.e. BioDare did not ask to ignore them), their goodness of fit was below 1 for FFT-NLLS and mFourfit or 0.9 for Spectrum Resampling and all estimates obtained by different methods were within 2.5h of each other. In Fig. 3, Figure 2 – supplement 1 and 2, the FFT-NLLS period estimates are shown. Period variability within and between cells was calculated as described previously (Kellogg & Tay, 2015).

Since some of the sections imaged overlap (e.g. Figure 2f; Figure 3a; Figure 2-figure supplement 1c, 2c), in order to not count cells multiple times, some of the cells were removed. This was done in the following manner: if there was an area of overlap in multiple sections, only cells belonging to the sections with lower *x* and *y* positions were kept, e.g. in Figure 3a, any cells in the upper hypocotyl section that also belong spatially to the lower hypocotyl section, were removed from subsequent analysis. In the repeat WT experiment the root tip section imaged encompasses a longer section of the root (Figure 2-figure supplement 1c). Hence, in order to make the analysis comparable to WT (Figure 3), we split the root tip section for further analysis. We considered the root tip cells of the repeat to be only those less than approximately 0. 66 mm from the actual tip, while the rest of them were classed as ‘Root up from tip’ (Figure 2-figure supplement 1a, b).

In the case of analysis at tissue level, where multiple sections had to be pooled for analysis (e.g. Figure 1g, h and Figure 1-figure supplement 3, 4, 5), since different sections were imaged at different times, before any further statistics were done, all the data was interpolated at the times where measurements across any section were made.

For analysis of amplitudes, peak and trough times for the individual cells (Figure 2a, f, g, and Figure 2-figure supplement 1a, e, f; Figure 2-figure supplement 2a, e, f) were identified using the findpeaks function in MATLAB. This was done on linearly detrended data. In case of the WT data (Figure 2), since the data is sampled more frequently (every 1.1h vs. 3h in WT repeat and CCA1-long line), the data is noisier, hence a smoothing filter (robust local regression using weighted linear least squares and a 2^nd^ degree polynomial model) was also applied after linear detrending. Amplitudes of traces were calculated as a mean of all trough to peak and peak to trough amplitudes.

### Luciferase space-time analysis

To facilitate analysis, individual seedlings were manually cropped into individual time stacks using ImageJ. From these image stacks a rectangular region of interest (ROI) containing the full length of the root and as much length of the hypocotyl as possible, whilst still excluding the cotyledons, was defined. Custom developed MATLAB scripts were used to extract luminescence data for each pixel in the ROI, giving time series for each pixel. After inspection of the images and the time series, some features were identified and the following measures applied to address them:

I. Occasionally the cotyledon of the seedling or of a neighboring seedling protrudes into the ROI. At this stage the ROI was checked for pixels of overlapping seedlings and these regions were manually removed from the affected frames.
II. Inside of the ROI the hypocotyl and root are surrounded by peripheral background pixels. The root and hypocotyl were segmented from the background using the mean of the grey levels as the threshold. The algorithm was applied to each image in the stack individually.
III. Commonly supposed to be from solar cosmic rays, pixel spikes in intensity values occur sporadically in images. A 3-by-3 pixel median filter is applied to each image to remove these spikes.
IV. The luminescence signal strength in a single seedling is weak and therefore the signal to noise ratio relatively low. A third order Butterworth filter was applied to pixel time series to remove high frequency noise. Time series were filtered using MATLAB’s filtfilt.m function, which performs in the forward and reverse direction to avoid phase distortion. A cut off frequency of 15% of the Nyquist frequency was identified as a best fit to our data.
V. In all experiments we observed dampening of the signal over time. Time series were therefore amplitude de-trended to better visualise spatial patterns. Time series were de-trended using the algorithms developed for the mFourfit toolkit (Edwards et al., 2010).

To visualise spatial patterns across the length of the root, space-time plots of the root luminescence were created (Fig. 3i, Figure 3-figure supplement 1). To do this we take the maximum signal intensity across one pixel wide longitudinal sections of the root for each image and assign this value to position *m,n* of the space time plot, where *m* is the image number and *n* the longitudinal section. The space-time plots presented include 10 pixels of the hypocotyl. The mean luminescence is normalised so that the peak expression of each longitudinal section (*n*) is 1.

### Model simulation

In Figure 1a we simulate an existing deterministic model of the clock (Pokhilko et al., 2012). The model was run for 168h from introduction into constant light conditions and *LHY/CCA1* mRNA is reported (in the model *CCA1* and *LHY* are treated as a single component (Pokhilko et al., 2012)).

In Figure 2, we simulated a stochastic model of an existing circadian clock model (Guerriero et al., 2012; Pokhilko et al., 2012). In Guerriero et al, the model is scaled by the parameter Ω, so that a molecule count close to Ω is obtained. For detailed description of the scaling, the reader can refer to this paper. Comparison of the model simulated for different Ω values to the previously published data indicates that the model molecule count of a few hundred cells (i.e. Ω) is a good prediction of the actual molecule count (Guerriero et al., 2012). Here we have taken the same circadian clock model and simulated it for various values of Ω. Model equations scaled for the Ω factor are given in (Guerriero et al., 2012). The model was simulated for 200h from introduction into constant light conditions and 100 simulation runs (proxy for 100 cells) were performed. The stochastic simulations were performed using the Gillespie algorithm (Gillespie, 1977). For each simulation, further analysis of amplitudes and period was done after the simulated data was interpolated at 2h intervals and then only for the simulated data from 28h to 168h in LL, in order to be closely comparable to the time interval of the original single cell data (Figure 1d). The Gillespie algorithm was written in MATLAB and the amplitudes and periods of the simulations were extracted using the MATLAB findpeaks function. Periods were calculated as a mean difference of peak-to-peak intervals. Amplitudes were calculated as a mean of all trough to peak and peak to trough amplitudes.

### Synchronisation analysis

For a set of individual cells, the inter-cellular synchrony was analysed. First, one cell was selected as a centroid of the synchronization analysis. Then, its neighboring cells, defined as those located within its sphere (radius *r*), were extracted. From CCA1-YFP expression signal, phase of the *j*-th neighboring cell (*j*=*1*,*2*..,*N*) was computed as (Pikovsky *et al.*, 2003)

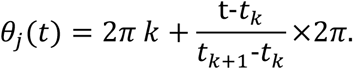

Here, the *k*-th peak time *t_k_* of the bioluminescence signal was detected by a cosine fitting method (coefficient of determination larger than 0.7) using the estimated period *τ_i_*. Then for each time point, the order parameter *R(t)* (Kuramoto, 1984) was obtained as

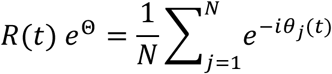

The order parameter (0<*R*<1) becomes unity for completely synchronized cells (θ_1_=θ_2_=..=θ_*N*_), whereas it becomes zero for non-synchronised cells. Figure 3-figure supplement 2 shows the results of synchronization analysis for root tip (**a**), lower root (**b**), upper root (**c**), lower hypocotyl (**d**), upper hypocotyl (**e**), and cotyledon (**f**). For each section, a total of *n* curves were drawn by selecting individual cells as the centroids (root tip, *n* = 242; lower root, *n* = 83; upper root, *n* = 46; lower hypocotyl, *n* = 114; upper hypocotyl, *n* = 53; cotyledon, *n* = 103). By linear regression analysis of each curve, the slope of the order parameter against time was computed, where a positive slope implies that the level of synchrony increases in time due to cell-to-cell interactions. Figure 3-figure supplement 2h shows the dependence of the slope value on cell density (ρ=N/(4/3)πr^3^). Positive slopes are mostly found in the root tip (Figure 3-figure supplement 2g), with a high correlation to the cell density.

Next, the coupling strength was estimated for each synchronization curve {*R*(*t*)}. Our approach is based upon a simplified version of the technique developed for weakly interacting mammalian circadian cells (Rougemont & Naef, 2007). As a model for the neighboring cells, we consider a set of coupled phase oscillators

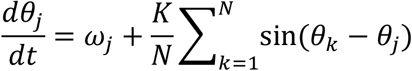

Assuming that the period τ_j_ estimated from the *j*-th cellular trace is not strongly affected by the other cells (Rougemont & Naef, 2007), the natural angular frequency was set as ω_j_=2π/τ_j_ for each oscillator. Given an initial condition θ_j_(0) extracted from the cellular traces, the phase oscillator model was simulated (Euler method with time step 0.1 h). Accordingly, the time evolution of the order parameter *R(t)* could be obtained. The coupling strength, which was initially set as *K*=0.002, is constant for each simulation. Staring from the minimum level of coupling, the coupling strength was slowly increased so that the phase oscillators are eventually mutually synchronized and the corresponding slope value increases monotonously. At the point when the slope value exceeds the one obtained from the experiment, the corresponding value of *K* provides the coupling estimate for the experimental data. Figure 3-figure supplement 2i shows the results. Stronger coupling was estimated for densely populated areas, implying that the cell-to-cell interactions are strengthened when cells are closely located to each other.

To examine the dependence of the present analysis on the synchrony measure used, the synchronization index (Garcia-Ojalvo, Elowitz, Strogatz, 2004) was utilized in place of the order parameter. The synchronization index has the advantage that the noise-sensitive procedure of phase extraction from the cellular traces is not required, since it can be computed directly from the measured signals. For *N* cellular traces {*x_j_*(*t*): *j*=1,2,..,*N*}, the averaged signal *M*(*t*) = (1/*N*) ∑_*j*_ *x*_*j*_(*t*) is computed. Then the synchronization index is given by

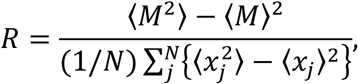

where <> denotes time average. In a synchronized cellular state, the averaged signal gives rise to a pronounced amplitude, resulting in *R*=1. The fully desynchronized cellular state, on the other hand, results in *R*=0. To see the time evolution of the level of synchrony, the synchronization index *R*(*t*) at time *t* was computed for windowed time traces of {*x*_*j*_,(*s*): *t*-12<*s*<*t*+12} (window: 24h). Figure 3-figure supplement 3 shows the analysis results based on the synchronization index. The panels are ordered in correspondence with those of Figure 3-figure supplement 2. Positive slopes are again found in the root tip, implying that the level of synchrony increases in time due to cell-cell interactions. The results are therefore consistent with the ones obtained by the order parameter.

To examine the dependence of the synchronization analysis on the particular experimental data set used, the order parameters were computed for the CCA1-long line. As shown in Figure 3-figure supplement 4, the slope of the order parameter for the CCA1-long line experiment is again well correlated with the cell density, where highly dense cells are located in the root tip. Although the time resolution was three times lower in this experiment, the same tendency was observed. The WT repeat experiment (Figure 2-figure supplement 1) was not analysed, as the timeseries was too short and time resolution was too low to enable accurate synchronization analysis.

### Phase oscillator model

We constructed a model where we describe the dynamics of the CCA1∷CCA1-YFP in each cell by a simple Kuramoto phase oscillator. For every cell at a position (*m,n*) in the plant (when viewed in 2D with *m* denoting position in the horizontal direction and *n* denoting position in the vertical direction) the phase of the oscillator *θ*^(*m,n*)^ changes in time (*t*) so that

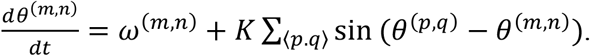

Here *ω*^(*m,n*)^ describes the intrinsic frequency of the oscillator and the second term describes the oscillator’s dependence on the coupling to the nearest neighbours (i.e. cells in positions (*p,q*) where *p*=*m*−1,..,*m*+1 and *q*=*n*−1,…,*n*+1) with *K* as the coupling constant. The bioluminescence of each cell is then taken to be 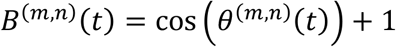. Cells in the plant conform to a template that is taken to be a symmetric shape, and resembles the shape of a seedling. The total bioluminescence across each vertical section (*n*) of the plant is taken to be the sum of the bioluminescence of all cells along that section i.e., 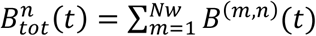 where the width of the plant counts *Nw* number of cells. The space-time plot shown in Figure 3k shows the total luminescence normalized so that the peak expression of each vertical section is 1.

In the model we assume that the cells in the three sections (the cotyledon/hypocotyl, the root and the root tip) have different intrinsic periods, with the cells in the cotyledon and hypocotyl having period of 24h, those in the root having the period of around 25.55h and the ones in the root tip a period of 22.67h. These overall match the qualitative period differences seen across the different plant sections. In all simulations the coupling constant *K* is arbitrarily set to 1. The ODEs are solved using the Euler method.

